# *Pseudomonas syringae* pv. *tomato* DC3000 induces defense responses in diverse maize inbred lines

**DOI:** 10.1101/2023.11.17.567553

**Authors:** Namrata Jaiswal, Matthew Helm

## Abstract

Many phytopathogens translocate virulence (effector) proteins into plant cells to circumvent host immune responses during infection. One such pathogen is *Pseudomonas syringae* pv. *tomato* DC3000, which secretes at least twenty-nine effectors into host cells, of which a subset elicits host defense responses in crop plant species. However, it is unknown whether *P. syringae* pv. *tomato* DC3000 activates immune responses in diverse maize inbreds. Here, we screened a diverse maize germplasm collection for effector-dependent recognition of this bacterial pathogen. As a control, we infiltrated *Pseudomonas syringae* DC3000(D36E), a derivative of *P. syringae* pv. *tomato* DC3000 that lacks all endogenous effectors. In our evaluations, we observed a variety of responses to *P. syringae* pv. *tomato* DC3000 in maize and scored the phenotypes as either no observable response (N) or as one of three responses: weak chlorosis (WC), chlorosis (C) with minimal cell death, and hypersensitive reaction (HR)-like cell death. Of the twenty-six maize inbreds screened, 13 were scored as N, 2 as WC, 2 as C, and 9 as HR-like cell death. Importantly, no maize line responded to *P. syringae* DC3000(D36E), demonstrating the responses observed are likely dependent upon recognition of one or more *Pseudomonas* effectors. Importantly, maize inbreds that recognize *P. syringae* pv. *tomato* DC3000 accumulated detectable hydrogen peroxide as well as an increase in transcript expression of a subset of maize defense genes. Collectively, our results will likely stimulate new research aimed at identifying the cognate maize disease resistance proteins that recognize the activities of one or more bacterial effectors.

## INTRODUCTION

To facilitate disease development, plant phytopathogens often secrete a repertoire of virulence proteins, known as effectors, to host cells during the infection process (Bentham et al., 2020). The primary function of these pathogen-secreted effectors is to manipulate host cell physiology and development, including the structure and function of plant cells as well as modulate host immune responses (Bentham et al., 2020; Lovelace et al., 2023). To circumvent the immune-suppressing activities of diverse plant phytopathogens, plants have evolved a two-tiered, integrated immune signaling network (Bentham et al., 2020; Lovelace et al., 2023; Ngou et al., 2022). The first tier, often referred to as cell surface-triggered immunity, primarily involves the detection of pathogen-associated molecular pattern (PAMPs) by cell surface-localized, transmembrane immune receptors (Bentham et al., 2020; Ngou et al., 2022). Upon recognition of pathogen-derived PAMPs, the host cell will initiate several coordinated defense responses including, in part, an increase in the production reactive oxygen species (ROS), stomatal closure, calcium influx and accumulation within the host cytosol, as well as up-regulation of defense-related genes (Bentham et al., 2020; Chen et al., 2022; Ngou et al., 2022). The second tier of the plant immune system is often mediated by intracellular innate immune receptors known as nucleotide-binding, leucine-rich repeat (NLR) proteins, whose primary function is to detect the activities of pathogen-secreted effectors inside host cells (Bentham et al., 2020; Ngou et al., 2022). Once activated, plant NLR proteins initiate a cell death response, known as the hypersensitive reaction (HR), which is thought to restrict pathogen growth and systemic spread (Bentham et al., 2020; Coll et al., 2011; Ngou et al., 2022). This tier of host immunity is often referred to as effector-triggered immunity (ETI) or intracellular receptor-mediated immunity (Ngou et al., 2022).

The bacterial phytopathogen *Pseudomonas syringae* pv. *tomato* DC3000 has been foundational to the study of effector biology and its study has yielded significant insights into the structure of the plant immune system (Wei and Collmer, 2018; Xin and He, 2013). During pathogenesis, this hemibiotrophic pathogen colonizes the host apoplast by entering through stomata and uses the type III secretion system (T3SS) to deliver at least twenty-nine effector proteins directly into the host cell cytoplasm (Wei et al., 2018; Xin and He, 2013). Once translocated, these effector proteins predominantly function to suppress host immune responses or modulate host development, thereby facilitating disease progression (Wei et al., 2018; Xin and He, 2013). For example, the *P. syringae* pv. *tomato* DC3000 effector HopM1 suppresses both PAMP-elicited ROS production as well as stomatal closure in Arabidopsis, thus facilitating disease development (Lozano-Duran et al., 2014). Additionally, the HopK1 effector from *P. syringae* pv. *tomato* DC3000 suppresses HR-like cell death elicited by another *P. syringae* pv. *tomato* DC3000 effector, HopA1, as well as PTI responses including ROS production and callose deposition (Guo et al., 2009; Jamir et al., 2004; Li et al., 2014). Interestingly, HopK1 encodes a chloroplast transit peptide and accumulates in host chloroplasts, suggesting chloroplast localization may be important for HopK1-mediated suppression of PTI and ETI responses (Li et al., 2014). HopQ1 (also known as HopQ1-1) is conserved across pathogenic bacteria and, intriguingly, induces an HR-like cell death response when transiently expressed in *N. benthamiana* (Ferrante et al., 2009; Wei et al., 2007). Upon *P. syringae* pv. *tomato* DC3000-dependent delivery into Arabidopsis, AvrPto and AvrPtoB function redundantly to attenuate flagellin22 (flg22) perception by the cell surface-localized immune receptor, FLS2 (Boller and Felix, 2009; Kvitko et al., 2009; Martin, 2012).

The maize Nested Association Mapping (NAM) recombinant inbred lines (RILs) representing global maize diversity were derived from twenty-six inbred founder lines and are a valuable genetic resource for discovering allelic diversity against plant pathogens of maize (Gage et al., 2020; Singh et al., 2023). For example, Kump and colleagues (2011) used the NAM population to identify thirty-two quantitative trait loci associated with resistance to *Cochliobolus heterostrophus*, the causative pathogen of Southern Leaf Blight disease, in maize. Furthermore, this mapping population was also used to identify the genetic resistance architecture underlying quantitative disease resistance to Gray Leaf Spot (GLS) disease and Northern leaf blight (NLB) disease (Benson et al., 2015; Poland et al., 2011). These studies thus demonstrate the value and versatility of screening the NAM population for identifying maize disease resistance genes. To the best of our knowledge, the maize NAM population has not been used to identify putative sources of genetic resistance to *P. syringae* pv. *tomato* DC3000.

In the current study, we screened the maize NAM population for recognition of the bacterial pathogen *P. syringae* pv. *tomato* DC3000. Of the twenty-six maize inbreds evaluated in our screen, 13 were scored as having no observable response to *P. syringae* pv. *tomato* DC3000. Two inbred lines were phenotyped as having a weak chlorotic response and two as having a chlorotic response with minimal cell death. The nine remaining maize inbreds all responded to *P. syringae* pv. *tomato* DC3000 with HR-like cell death. Consistent with the phenotypic responses, the maize inbred lines that recognized *P. syringae* pv. *tomato* DC3000 accumulated detectable hydrogen peroxide within the infiltrated regions as well as an increase in transcript expression of a subset of maize defense genes. As a control, we infiltrated an effectorless mutant strain of *Pseudomonas syringae* DC3000, termed *P. syringae* DC3000(D36E), which did not elicit observable defense responses or hydrogen peroxide accumulation in any maize inbred line. These data thus suggest *P. syringae* induces defense responses in diverse maize inbreds and that maize likely encodes disease resistance proteins that recognize the activities of one or more *Pseudomonas* effectors.

## MATERIALS AND METHODS

### Plant growth conditions

Seed of maize inbreds were ordered from the North Central Regional Plant Introduction Station via the National Plant Germplasm System Web Portal. Maize inbreds were grown in plastic pots containing Berger Seed Germination and Propagation Mix supplemented with Osmocote slow-release fertilizer (14-14-14). Plants were maintained in a growth chamber with 16h:8h photoperiod, light:dark at 24°C with light and 22°C in the dark and 60% humidity with average light intensities at plant height of 130µmols/m^2^/s.

### *Pseudomonas* growth conditions and leaf infiltration assay

*Pseudomonas syringae* pv. *tomato* DC3000 and *Pseudomonas syringae* DC3000(D36E) strains were grown on King’s B (KB) medium supplemented with 25µg of rifampicin and 50µg of kanamycin per milliliter for two days at 30°C. The *Pseudomonas* strains were resuspended in 10mM MgCl_2_ to an optical density at 600 nm (OD_600_) of 0.5. Bacterial suspensions were infiltrated into the abaxial surface of the second leaf of two-week old maize seedlings using a 1-mL disposable syringe, and the infiltrated regions were indicated with a black permanent marker. The leaf surface was nicked with a sterile razor blade prior to infiltration to facilitate fluid entry. The infiltrated leaves were phenotyped five days following *Pseudomonas* infiltration and a representative leaf was photographed under white light shortly thereafter.

### Histochemical detection of hydrogen peroxide accumulation

Hydrogen peroxide (one of several reactive oxygen species) accumulation was detected as previously described with slight modifications (Bach-Pages and Preston, 2017). Briefly, the *Pseudomonas* suspensions were infiltrated into two-week old maize seedlings and, 3 days post-infiltration, leaf segments were harvested and incubated in freshly prepared 3,3’-diaminobenzidine (DAB) solution (1 mg/mL (Sigma-Aldrich) dissolved in distilled water (pH 3.8) overnight with gentle agitation in the dark. After overnight incubation, the DAB-stained leaf tissues were rinsed with distilled water, submerged in absolute ethanol, and incubated at 65°C for 2 hours to clear the chlorophyll. The leaf segments were rehydrated with 75% ethanol and incubated for 1 hour at room temperature, followed by incubation with 50% ethanol until the leaves were cleared of chlorophyll. The cleared leaf tissue was stored in ultrapure water (supplemented with 20% glycerol) for photography. Representative DAB-stained leaf segments were photographed under white light.

### RNA extraction and reverse transcriptional real-time quantitative PCR of maize defense genes

Two-week old maize leaves (cvs. B73 [non-responder] and Mo18W [responder]) were infiltrated with *P. syringae* pv. *tomato* DC3000. The infiltrated leaf tissue was harvested 12- and 24-hours post-*Pseudomonas* infiltration, flash frozen in liquid nitrogen, and stored at -80°C until processing. Total RNA was isolated using TRIzol reagent from maize tissues according to the manufacturer’s instruction (Direct-zol^TM^ RNA miniprep, ZymoResearch) followed by in-column DNase I treatment. RNA concentration was measured using Nanodrop (Thermo Scientific). Two micrograms of total RNA were used for first-strand cDNA synthesis using AMV reverse transcriptase (New England Biolabs). A quantitative real-time PCR (RT-qPCR) assay was performed on a LightCycler 480 II system using 2x SYBR Green qPCR master mix (BioRad). Maize GAPDH (Zm00001eb233140) was chosen as an endogenous reference gene to normalize the data. Three technical replicates of the RT-qPCR assay were used for each sample with a minimum of three biological replicates. Each biological replicate was sampled from three individual plants. The transcript expression level was calculated using the 2^-ΔΔCT^ method (Livak and Schmittgen, 2001). The experiment was repeated at least two independent times. Primers used for RT-qPCR are as follows (5’ to 3’): ZmGAPDH-FW (CCTGCTTCTCATGGATGGTT); ZmGAPDH-RV (TGGTAGCAGGAAGGGAAACA); ZmPR3-FW (AGCTTCCTCGTCTCATCATCG); ZmPR3-RV (TGTTGACGTAGGCGTAGAGC); ZmPR5-FW (GCAGCCAGGACTTCTACGAC); ZmPR5-RV (CACAGGCATGGGTCTTCAC); ZmPAL-FW (TCTGGTCCCGCTCTCCTACA); ZmPAL-RV (TTGAAGAAGCCGCCCTCGAT); Peroxidase-FW (CTCATCAACCACCCGGACAC); Peroxidase-RV (TGCAGCAAGCCTTATTTGAACA).

## RESULTS

The observation that *P. syringae* pv. *tomato* DC3000 elicits robust immune responses in crop plant species such as wheat led us to hypothesize that this bacterial phytopathogen may also activate defense responses in maize inbreds. To test our hypothesis, we infiltrated the parental inbred lines of the maize nested association mapping population with *P. syringae* pv. *tomato* DC3000 and phenotyped their responses five days post-*Pseudomonas* infiltration (Figure 1A). As a control, we infiltrated *P. syringae* pv. *tomato* DC3000 into wheat leaves, which was previously shown to recognize this bacterial pathogen (Ramachandran et al., 2017). In our evaluations, we observed a variety of responses to *P. syringae* pv. *tomato* DC3000 and scored the phenotypes as either no observable response (N) or as one of three responses: weak chlorosis (WC), chlorosis (C) with minimal cell death, and hypersensitive reaction (HR)-like cell death. Representative examples of phenotypic responses are shown in Figure 1B, and a complete list of inbred lines and their responses are shown in Table 1. Of the twenty-six maize inbreds tested, 13 were scored as N, 2 as WC, and 2 as C (Figure 1B; Table 1). Nine maize inbred lines (CML52, CML69, CML247, CML333, HP301, M162W, Mo18W, Ms71, and IL14H) displayed a HR-like response to *P. syringae* pv. *tomato* DC3000 within the infiltrated region (Figure 1B; Table 1). As a control, we infiltrated a mutant strain of *P. syringae* pv. *tomato* DC3000, termed *P. syringae* DC3000(D36E), which lacks all endogenous bacterial effectors (Carter et al., 2019; Wei et al., 2015). As expected, there was no observable response in any inbred line tested to *P. syringae* DC3000(D36E) (Figure 1B). These data thus demonstrate the phenotypic responses observed in response to *P. syringae* pv. *tomato* DC3000 are likely dependent upon recognition of one or more *P. syringae* effectors.

**Figure 1.**
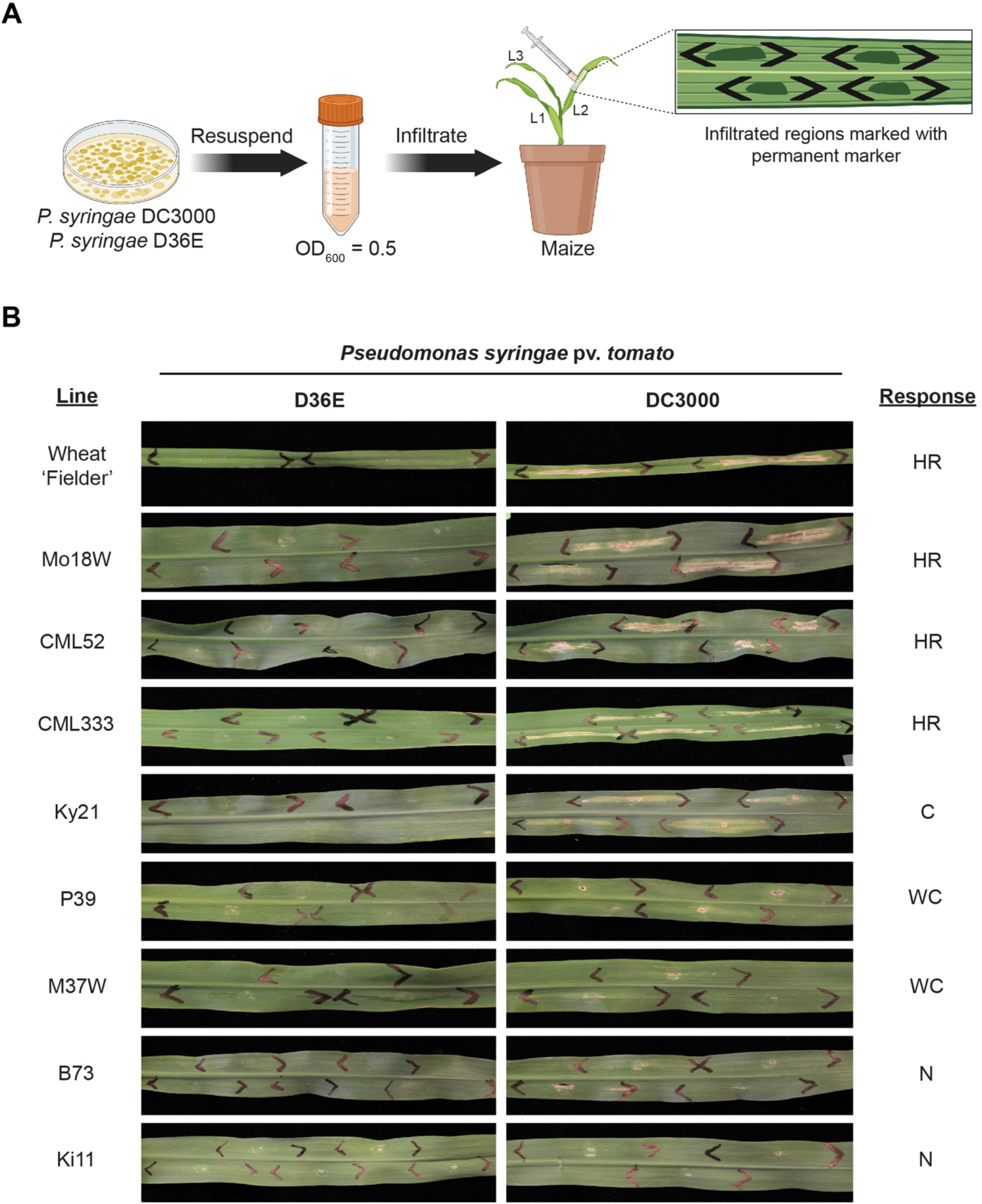
*Pseudomonas syringae* pv. *tomato* DC3000 induces defense responses in maize. **A)** Schematic diagram of the *Pseudomonas* HR assay. The *P. syringae* pv. *tomato* DC3000 and *P. syringae* DC3000(D36E) strains were resuspended in 10mM MgCl_2_. The bacterial suspensions were adjusted to an optical density at 600nm (OD_600_) of 0.5 and syringe-infiltrated into the underside surface of the second leaf (L2) of two-week old maize seedlings. The infiltrated regions were marked with a black permanent marker. **B)** Responses of the maize inbred lines to either *P. syringae* pv. *tomato* DC3000 or *P. syringae* DC3000(D36E) were phenotyped five days following infiltration. Responses were scored as N, no observable response; WC, weak chlorosis; C, chlorosis with cell death; and HR, hypersensitive reaction. Wheat cv. Fielder inoculated with either *P. syringae* pv. *tomato* DC3000 or *P. syringae* DC3000(D36E) was used as a control (Carter et al., 2019; Ramachandran et al., 2017).

**Table 1.**
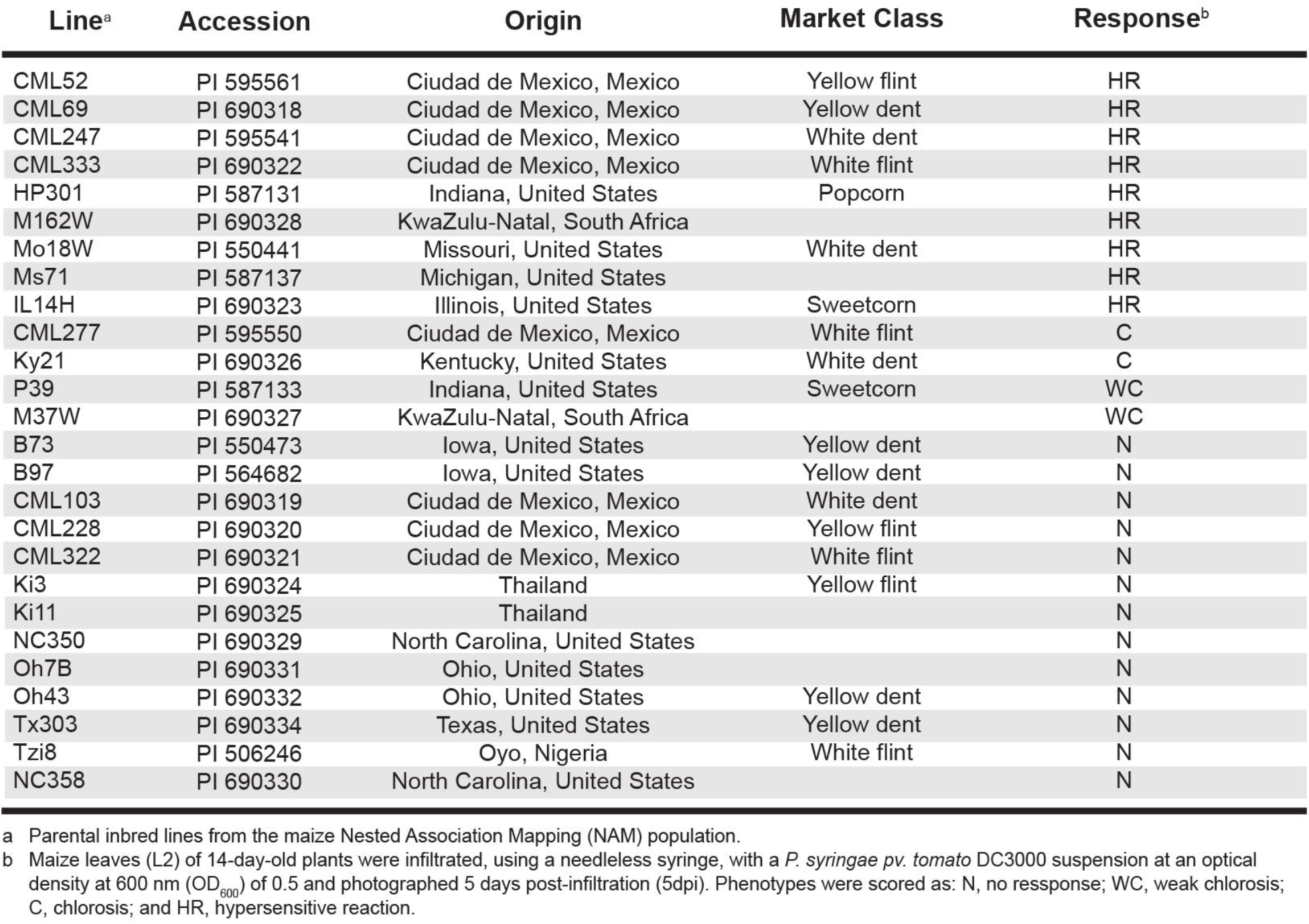
Responses of the twenty-six inbred maize lines of the nested association mapping (NAM) to *Pseudomonas syringae* pv. *tomato* DC3000. Accession numbers, origins, and market class were mined from the Maize Genetics and Genomics Database.

The production and accumulation of reaction oxygen species, including hydrogen peroxide, is often used as a proxy for activation of defense responses (Carter et al., 2019). Therefore, we used 3,3’-diaminobenzidine (DAB) staining following leaf infiltration with either *P. syringae* pv. *tomato* DC3000 or *P. syringae* DC3000(D36E) to test whether the phenotypic responses of some maize inbred lines to *P. syringae* pv. *tomato* DC3000 is correlated with hydrogen peroxide accumulation. Consistent with the phenotypic responses, maize inbreds that displayed an HR-like cell death response to *P. syringae* pv. *tomato* DC3000 accumulated detectable hydrogen peroxide within the infiltrated region (Figure 2). In contrast, there was no hydrogen peroxide detected in maize lines that did not recognize *P. syringae* pv. *tomato* DC3000 or in lines infiltrated with *P. syringae* DC3000(D36E) (Figure 2). These results thus reveal the chlorotic and HR-like cell death response in maize inbreds that recognize *P. syringae* pv. *tomato* DC3000 is strongly correlated with the production and accumulation of ROS.

**Figure 2:**
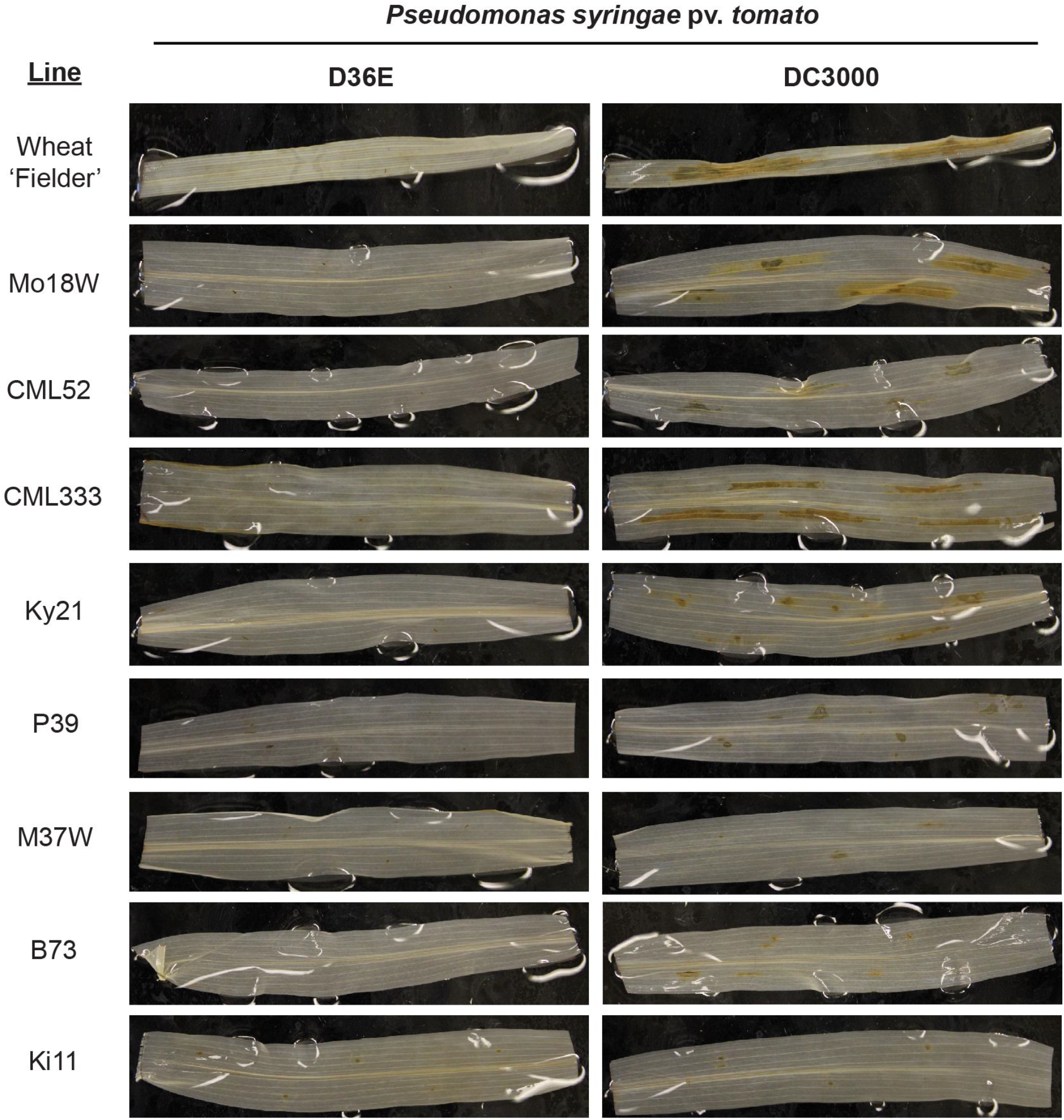
Maize lines accumulated detectable hydrogen peroxide in response to *P. syringae* pv. *tomato* DC3000. Representative maize inbreds assayed and *Pseudomonas* treatments were as in Figure 1B. Three days following *Pseudomonas* infiltration, leaf segments were harvested from the infiltrated regions and stained overnight with 3,3’-diaminobenzidine (DAB) solution. After overnight incubation, the leaf tissue was rinsed with deionized water, cleared with ethanol to remove the chlorophyll, and stored in ultrapure water (supplemented with 20% glycerol) for photography. Wheat cv. Fielder inoculated with either *P. syringae* pv. *tomato* DC3000 or *P. syringae* DC3000(D36E) was used as a control.

Though chlorotic and cell death responses are considered defense responses in grasses (Carter et al., 2019; Smith and Mansfield, 1981; Yin and Hulbert, 2010), it was unclear whether the *P. syringae* pv. *tomato* DC3000-mediated defense responses in maize were correlated with increased defense gene expression. We, therefore, used reverse transcription-quantitative polymerase chain reaction (RT-qPCR) to measure mRNA abundance of a subset of defense-related maize genes in both non-responding (B73) and a responding (Mo18W) maize inbred lines at 12- and 24-hours post-*Pseudomonas* infiltration. In our analysis, we selected a subset of defense-related maize genes whose transcriptional activation is associated with salicylic acid signaling and which have been previously shown to be up-regulated during maize-pathogen interactions (Przybylska et al., 2018; Shumilaket al., 2023). Consistent with the phenotypic responses to *P. syringae* pv. *tomato* DC3000, transcript expression of the selected defense genes was significantly upregulated in Mo18W when compared to B73 at 12- and 24-hours following *Pseudomonas* infiltration (Figure 3). Collectively, these data clearly demonstrate *P. syringae* pv. *tomato* DC3000 activates robust defense responses, including hydrogen peroxide accumulation as well as up-regulation of defense-related genes, in diverse maize inbreds.

**Figure 3:**
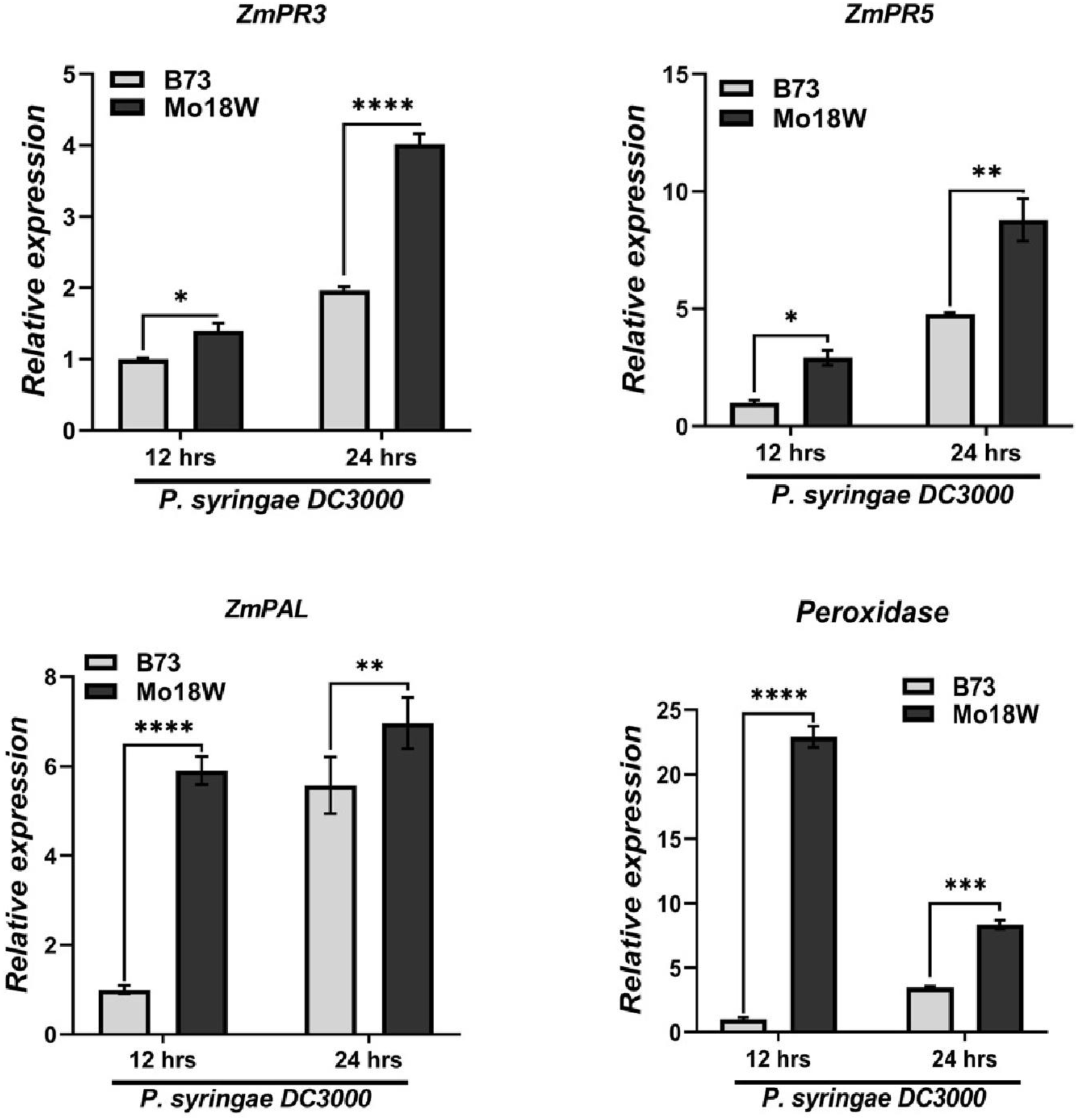
*P. syringae* pv. *tomato* DC3000 significantly increases transcript expression of maize defense genes. Reverse transcription-quantitative polymerase chain reaction (RT-qPCR) expression analyses of selected maize defense genes: Pathogenesis-related 3 (ZmPR3), Pathogenesis-related 5 (ZmPR5), Phenylalanine Ammonia Lyase (ZmPAL), and peroxidase in response to *P. syringae* pv. *tomato* DC3000. Two-week old maize leaves (cvs. B73 [non-responder] and Mo18W [responder]) were infiltrated with *P. syringae* pv. *tomato* DC3000. The infiltrated leaf tissue was harvested 12- and 24-hours post-*Pseudomonas* infiltration, flash frozen in liquid nitrogen, and stored at -80°C until processing. Total RNA extraction followed by cDNA preparation and expression of defense genes was quantified by RT-qPCR. Values represent mean ± S.D. (n = 3) from three biological replicates and three technical replicates in each experiment. The maize GAPDH gene was used as an internal reference gene. The gene expression in B73 plants at 12 hours after *P. syringae* pv. *tomato* DC3000 infiltration was set to 1. Multiple comparisons were conducted by two-way ANOVA followed by the Šídák’s method. One asterisk (*) indicates P<0.05, two (**) P<0.01, three (***) P<0.001, and four (****) P<0.0001.

## DISCUSSION

Our work presented here is, to the best of our knowledge, the first to show that *P. syringae* pv. *tomato* DC3000 elicits HR-like cell death in some maize inbreds. Consistent with our observations, this bacterial pathogen also induces defense responses in both *Nicotiana benthamiana* and soybean via the recognition of the HopQ1-1 and AvrD effectors, respectively (Kobayashi et al., 1989; Wei et al., 2007). Though our work shows that maize recognizes *P. syringae* pv. *tomato* DC3000, it remains unknown which *Pseudomonas* effectors are specifically recognized. To this end, Ruiz-Bedoya and colleagues (2023) recently generated thirty-six coisogenic *Pseudomonas* strains, each expressing a single *Pseudomonas* effector in the effectorless *P. syringae* DC3000(D36E) mutant. Hence, future research efforts will focus on infiltrating the library of *P. syringae* DC3000(D36E) coisogenic strains in maize to determine which bacterial effectors are specifically recognized (Ruiz-Bedoya et al., 2023). Such knowledge will be valuable in determining the molecular and cellular mechanisms underlying nucleotide-binding leucine-rich repeat immune receptors (NLR)-dependent recognition of *P. syringae* pv. *tomato* DC3000 effectors (Prigozhin et al., 2022; Thatcher et al., 2023).

The observation that *P. syringae* pv. *tomato* DC3000 elicits defense responses in maize suggests this bacterial pathogen can be utilized as a surrogate for expression of crop pathogen effectors in maize. Indeed, the bacterial type III secretion system from *P. syringae* pv. *tomato* DC3000 has been effectively used to deliver candidate effector proteins directly inside host cells to ascertain effector functions (Fabro et al., 2011; Qi et al., 2018; Ramachandran et al., 2017; Sohn et al., 2007). Development of a *P. syringae* pv. *tomato* DC3000-based heterologous expression system in maize will thus likely facilitate the functional screening of effector proteins from obligate biotrophic pathogens of maize. For example, effector proteins from the maize fungal pathogens *Ustilago maydis* and *Puccinia polyspora* suppress PAMP-elicited reactive oxygen species production (Chen et al., 2022; Navarrete et al., 2021; Saado et al., 2022). However, it remains unknown whether these crop pathogen effectors are capable of suppressing *P. syringae* pv. *tomato* DC3000-mediated defense responses in maize.

In summary, we show that *P. syringae* pv. *tomato* DC3000, but not the effectorless mutant, induces robust defense responses in some maize inbreds. Though it is unclear which *P. syringae* pv. *tomato* DC3000 effectors are recognized, our data nevertheless suggests maize likely encodes disease resistance proteins that recognize the activities of one or more *Pseudomonas* effectors. Future research efforts will, in part, focus on fine mapping the genetic loci responsible for *P. syringae* pv. *tomato* DC3000 recognition.

## ACKNOWLEDGEMENTS

We thank the North Central Regional Plant Introduction Station for providing seed of the maize inbred lines and Dr. Terri Cameron (USDA-ARS) for their technical assistance and maintenance of maize germplasm.

